# From GWAS to signal validation: An approach for estimating genetic effects while preserving genomic context

**DOI:** 10.1101/2023.03.09.531909

**Authors:** Scott Wolf, Varada Abhyankar, Diogo Melo, Julien F. Ayroles, Luisa F. Pallares

## Abstract

Validating associations between genotypic and phenotypic variation remains a challenge, despite advancements in association studies. Common approaches for signal validation rely on gene-level perturbations, such as loss-of-function mutations or RNAi, which test the effect of genetic modifications usually not observed in nature. CRISPR-based methods can validate associations at the SNP level, but have significant drawbacks, including resulting off-target effects and being both time-consuming and expensive. Both approaches usually modify the genome of a single genetic background, limiting the generalizability of experiments. To address these challenges, we present a simple, low-cost experimental scheme for validating genetic associations at the SNP level in outbred populations. The approach involves genotyping live outbred individuals at a focal SNP, crossing homozygous individuals with the same genotype at that locus, and contrasting phenotypes across resulting synthetic outbred populations. We tested this method in *Drosophila melanogaster*, measuring the longevity effects of a polymorphism at a naturally-segregating cis-eQTL for the *midway* gene. Our results demonstrate the utility of this method in SNP-level validation of naturally occurring genetic variation regulating complex traits. This method provides a bridge between the statistical discovery of genotype-phenotype associations and their validation in the natural context of heterogeneous genomic contexts.

## Introduction

Understanding how genetic variation regulates phenotypic differences between individuals is one of the main challenges of modern biology. Advances in genetic mapping methods, such as GWAS, have identified thousands of genetic variants associated with variation in complex traits across a wide range of organisms (Alsheikh et al., 2022; Pallares, Lea, et al., 2023; Saul et al., 2019; Visscher et al., 2017). Although mapping studies have contributed substantially to elucidating the genetic architecture of complex traits, the validation of candidate variants has dramatically lagged behind (Gallagher & Chen-Plotkin, 2018) and remains particularly difficult.

Methods for validating genetic association signals fall under two broad categories, methods that operate at the gene level and those that operate at the polymorphism level. Currently, the most common approaches to validate candidate genotype-phenotype associations rely on gene level experimental perturbation using loss-of-function mutations or RNAi constructs, comparing organisms with and without a functional copy of a gene of interest and assessing the phenotypic consequences (Bellen et al., 2019; Housden et al., 2017; Zimmer et al., 2019). Despite their immense usefulness in defining gene function, these approaches have important limitations when validating genotype-phenotype associations given how disconnected the validation context is from the association study that generated the signal of interest. In addition, the genetic effects assessed with such approaches usually represent genetic variation (e.g., null KO alleles) that does not segregate in natural populations. Recent developments in CRISPR technology provide an alternative that has revolutionized the field.

They have allowed for more realistic experiments in which specific single nucleotides can be targeted and replaced to assess the phenotypic effects of alternative variants (Hoedjes et al., 2023; Ramaekers et al., 2019). These CRISPR-based validation methods still present major drawbacks. First, off-target effects can be substantial and difficult to assess (Lessard et al., 2017; Schaefer et al., 2017). Secondly, CRISPR assays can be costly, time-consuming, and not readily available depending on the target organism. Critically, both loss-of-function and CRISPR-based studies usually modify the genome of a single genetic background, often using inbred lines (Mokashi et al., 2021). We argue that, given that genetic effects are often background-dependent (Chandler et al., 2013), working on a single genetic background limits the inference and generalizability of the validation results.

To address these challenges, we propose a simple, low-cost experimental scheme for the validation of genetic association at the polymorphism level in outbred populations, thus preserving variation in the genetic background across individual. The approach relies on three steps: 1) genotyping live outbred individuals at a focal SNP, 2) crossing homozygous individuals that share the same genotype at the locus of interest, and maintaining them as outbred populations, and 3) measuring and comparing phenotypes of interest across the resulting synthetic outbred populations that are fixed for the polymorphism of interest but have randomized genetic backgrounds (fig. 1).

**Figure 1:**
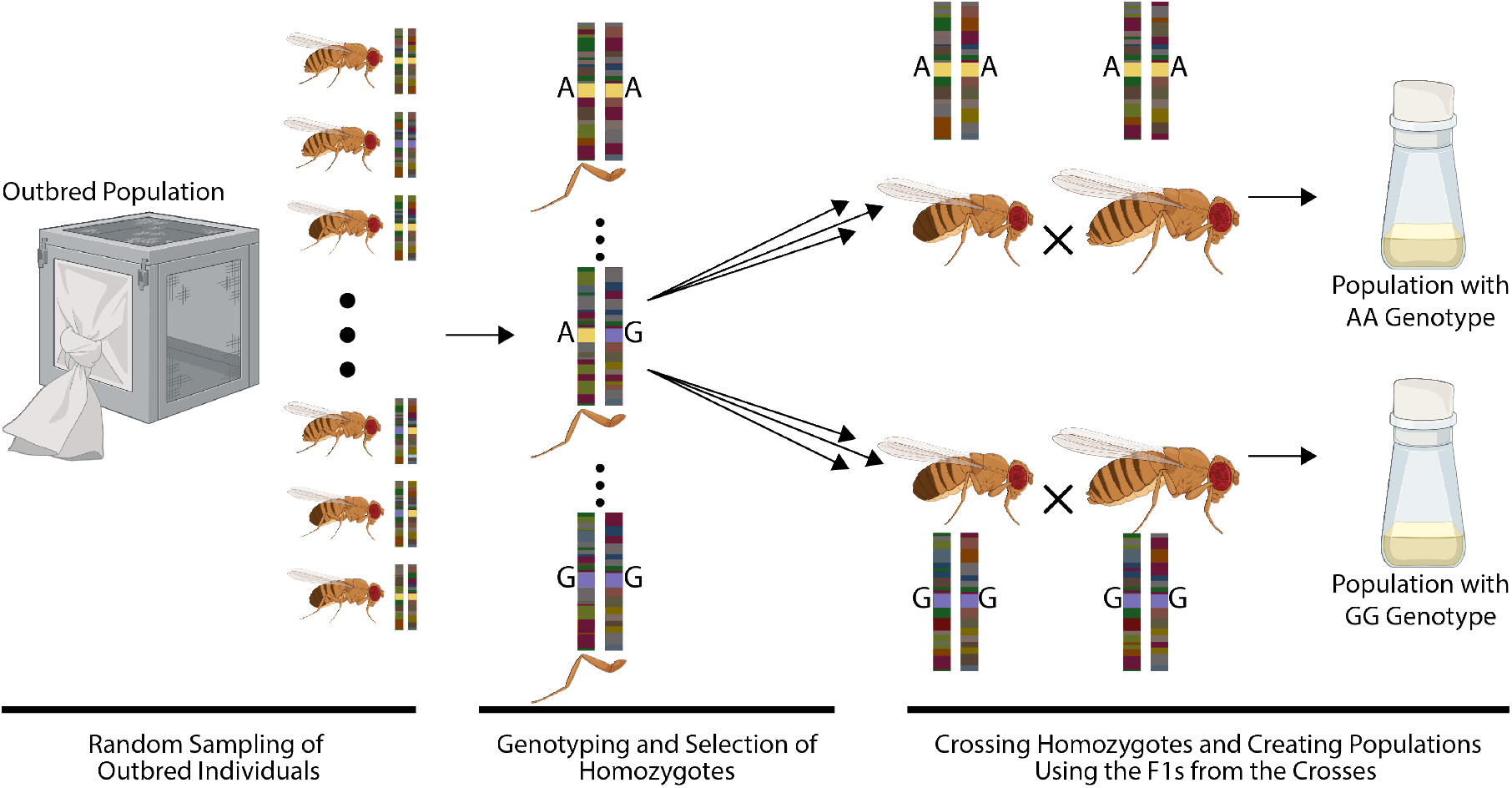
Experimental approach used to create populations with diverse genetic backgrounds fixed at a focal SNP. Virgin flies from both sexes are collected from an outbred fly population harboring the polymorphisms of interest. While anesthetized with CO2, a single leg is removed from each virgin fly, and individuals are placed into separate vials waiting for the legs to be genotyped. DNA is extracted from each leg and genotyped individually at the focal SNP using PCR and amplicon sequencing. Once males and females homozygous for the same genotype are identified, they are crossed (within genotype) and an equal number of offspring from each cross are transferred to a bottle to ensure that the genomic background of each founder individual is represented in the resulting population. In each bottle, all individuals are fixed for a given genotype at the locus of interest and the populations maintain genetic diversity present in the initial outbred population. The resulting populations are ready to be phenotyped for any trait of interest.

We demonstrate the utility of this approach in *Drosophila melanogaster* focusing on the phenotypic effects of a cis-regulatory polymorphism for *midway*. The *midway* gene is known in *Drosophila* to be involved in lipid metabolism (Buszczak et al., 2002; Girard et al., 2021; Tian et al., 2011), immune function (Tschapalda et al., 2016), and female fertility (Schüpbach & Wieschaus, 1991). And, it was recently identified as a lifespan-regulating gene in a study investigating the genetic basis for variation in lifespan in *D. melanogaster* (Pallares, Lea, et al., 2023). Furthermore, the mammalian ortholog of *midway, DGAT1*, has previously been linked with longevity in mice (Streeper et al., 2012). Pallares, Lea, et al. (2023) identified that *midway* plays a role in regulating lifespan using a loss-of-function mutation in an inbred line, we do not know if or how naturally-segregating genetic variants linked to *midway* expression are indeed responsible for lifespan variation in this species. Simultaneously, another study aimed at mapping genetic variants that regulate variation in gene expression levels genome-wide (i.e. eQTLs) in an outbred fly population, identified a candidate cis-eQTL upstream of *midway* (Pallares, Melo, et al., 2023). We used the validation paradigm outlined above to determine whether this regulatory variant identified for *midway* contributes to variation in lifespan.

To accomplish this, we created two fly populations, each homozygous for one of the eQTL focal alleles (i.e. AA vs GG) and, for each, quantified variation in longevity. We were able to validate the role this specific eQTL variant plays in lifespan and indirectly confirm that transcriptional variation in *midway* drives its effect on lifespan. Our experimental approach allowed us to validate a statistical discovery across a randomized set of genetic backgrounds in an outbred population. This study is one of the few validations of naturally occurring genetic variation controlling complex traits at the single nucleotide level.

## Methods and Results

### Drosophila melanogaster outbred populations

The candidate SNP for the gene *midway* we are investigating was identified in an eQTL mapping study that used an outbred mapping population of *D. melanogaster* derived from the Netherlands (NEX) (Pallares, Melo, et al., 2023). This population was generated by crossing 15 inbred Global Diversity Lines (Grenier et al., 2015), followed by ∼130 generations of recombination. The identification and initial validation of *midway* as a regulator of *Drosophila* lifespan was recently published as part of a GWAS-like study that used an outbred population derived from 600 isofemale lines caught in Princeton, NJ (Pallares, Lea, et al., 2023). To validate the effect of the *midway* eQTL on longevity, we used the same outbred NEX population where the eQTL was initially discovered. Flies were maintained at 25 °C, 65% relative humidity, and a 12h:12h light:dark cycle, and were fed media with the following composition: 1% agar, 8.3% glucose, 8.3% yeast, 0.41% phosphoric acid (7%), and 0.41% propionic acid (83.6%).

### Experimental populations used for SNP validation

To evaluate the effect of the target SNP on longevity, we followed the scheme described in fig. 1 and created two synthetic outbred populations with hundreds of individuals homozygous for each *midway* cis-eQTL allele (AA or GG) identified in (Pallares, Melo, et al., 2023). The candidate SNP eQTL for *midway* is located at 2L:16,812,901 (3759 bp upstream of the *midway* gene) and has a minor allele frequency of 16% in the NEX population where it was discovered (Pallares, Melo, et al., 2023).

We randomly selected virgin male and female flies from the outbred NEX population to identify flies homozygous at the focal locus. While the flies were anesthetized with CO2, we removed one leg from each individual for DNA extraction and genotyping (removing a leg does not alter viability). The flies were kept in separate vials until their genotypes were determined, after which they were paired with flies of the same genotype and mated. DNA was isolated with the QuickExtract™ DNA Extraction (cat no. QE09050) and the region around the focal *midway* polymorphism was amplified using the following primers: Fwd-TCGTCGGCAGCGTCAGATGTGTATAAGAGACAGGAGGAGCCACCAAGTGTTGT and Rev-GTCTCGTGGGCTCGGAGATGTGTATAAGAGACAGATCGAACTTCTCTCGCGACT. The sequencing library for these amplicons was generated using Illumina i5 and i7 primers. After bead cleaning, the library was sequenced using a MiSeq Nano flow cell (150bp PE reads) at the Genomics Core Facility of Princeton University. Genotypes were called using bcftools mpileup (Li, 2011) with parameters -Ou -B -q 60 -Q 28 -d 1000 -T -b files | bcftools call -Ou -m -o. After identifying the homozygous individuals for the two alleles, five crosses were set up in vials, each with one male and one female of the same genotype. From each cross of the same genotype, 10 male and 10 female offspring were combined in a bottle and allowed to mate freely for two generations before starting the survival assay. This design ensures that the genotype of each founding parent is represented in the synthetic outbred population (fig. 1).

To assess whether this candidate *midway* variant is associated with variation in lifespan, we performed survival assays in both experimental populations (i.e. one for each alternative genotype). One hundred males and one hundred females (1-2 days old) for each homozygous *midway* cis-eQTL genotype were distributed across 10 vials with 10 individuals of each sex in each vial. Flies were transferred onto fresh media every 3 days, and survival was monitored each time flies were transferred until the last fly died on day 79 (fig. 2).

**Figure 2:**
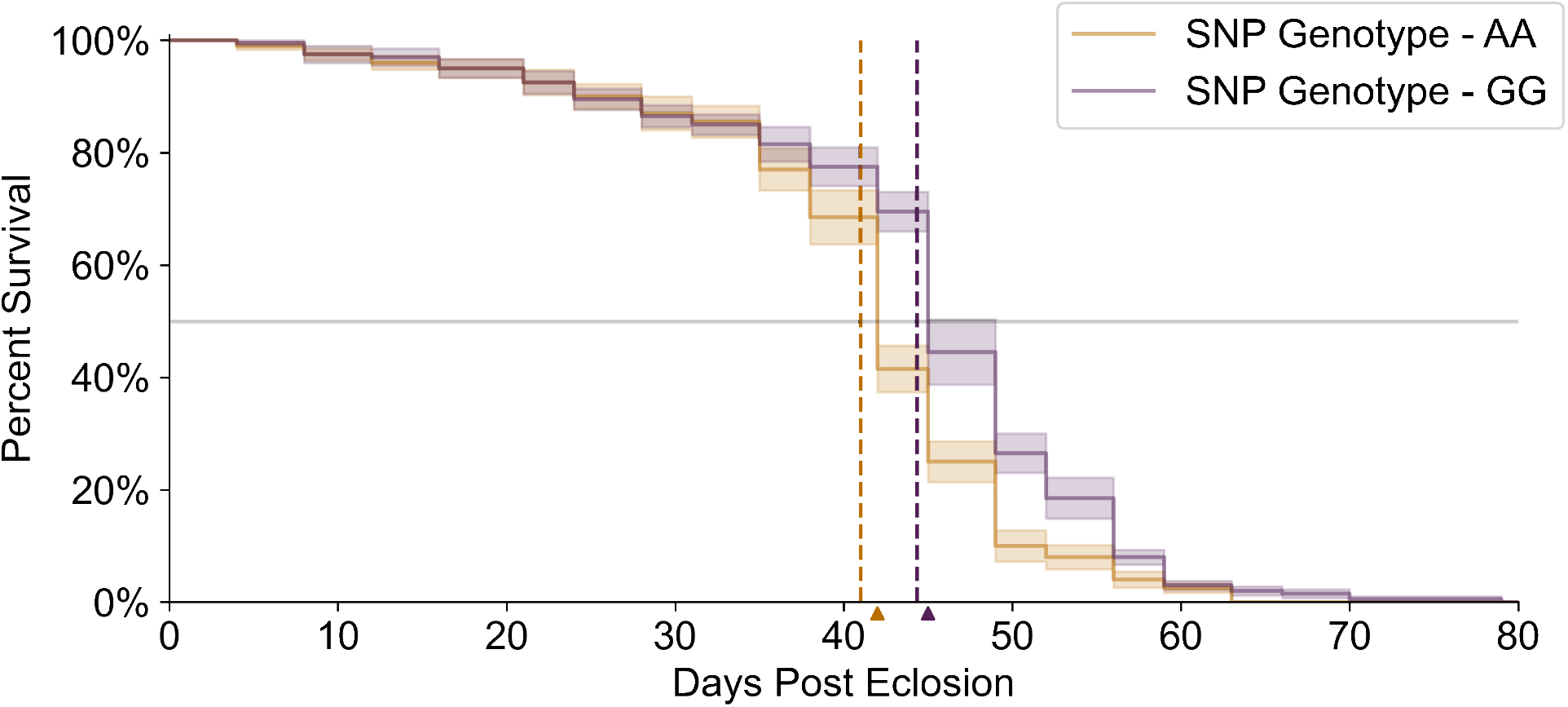
Effect of the *midway* regulatory polymorphism on *D. melanogaster* survival. Survival distribution stratified across midway genotype. Shaded regions show standard error, and the dashed vertical lines show mean lifespan by genotype (AA mean lifespan 40.99 days, GG mean lifespan 44.33 days). T50, the last day when 50% of individuals are alive, is denoted by wedges along the x-axis. T50 is day 42 for genotype AA and day 45 for genotype GG. Survival data for 200 individuals from each midway cis-eQTL genotype is included.

We performed a proportional hazard regression using the empirical survival distribution using the statsmodels Python package (Cox, 1972; Seabold & Perktold, 2010). Given that male and female *D. melanogaster* differ in average lifespan (Austad & Fischer, 2016), we were interested in whether the focal genotype modulated this difference. Therefore, our model was initially specified as:

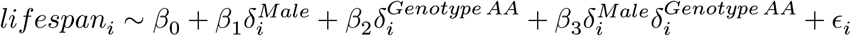

We did not find significant genotype-by-sex effects (95% CI 0.6635 -1.4597; p = 0.9365). We then proceeded with the following model:

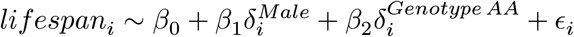

Our results show that after accounting for sex, flies homozygous for the minor GG genotype live longer than flies homozygotes for the major allele AA genotype. Cox’s proportional hazard regression using the final model yields a hazard ratio for genotype AA of 0.6767 (95% CI 0.5544 -0.8261; p = 0.0001).

## Discussion

The fast-growing power of mapping studies has produced an ever-expanding list of candidate genetic variants associated with complex trait variation. This large number of candidates with small effect sizes presents a formidable challenge for validating these signals, especially if the effects are context-dependent. To overcome some of these challenges, we developed an alternative experimental approach for candidate SNP validation that does not rely on genetic engineering (e.g. CRISPR or RNAi). Our approach relies on generating populations of individuals fixed for alternative genotypes of a target polymorphism but randomized across diverse genetic backgrounds. These synthetic outbred populations enable the validation or estimation of additive allelic effects of variants in a natural genetic context. This approach is facilitated by the low cost and high throughput of amplicon sequencing using Illumina’s MiSeq sequencer (i.e. hundreds of individuals can be genotyped-by-sequencing in a couple of days and at a low cost).

One of the key benefits of this approach is the gain in statistical power provided by the change in allele frequency between alternative genotypes in the synthetic outbred populations. While in the initial mapping population, potentially low minor allele frequency limits statistical power to detect variation in allelic effects, our approach yields a validation population where jointly, the minor allele frequency is 0.5 (e.g. in this study, the frequency of the minor allele in the mapping population was 0.16 and effectively rose to 0.5 across our validation populations). Thus, this approach maximizes the chance of identifying a phenotypic effect associated with a focal SNP of interest.

An additional benefit of having outbred populations fixed at alternative genotypes is that one can interrogate the pleiotropic effects of a given polymorphism by simply phenotyping each population for any number of traits and without any additional genotyping costs. For example, one could simultaneously and robustly estimate the effect of a SNP on gene expression and its effect on higher-order phenotypes such as behavior. This method can also be directly extended to assess the effect of candidate SNPs across a variety of environmental contexts and treatments, which opens up a wide range of possibilities, such as validating candidate genotype-by-environment interactions.

Here, we have connected two independent discoveries, derived from two genome-wide analyses in different populations of outbred *D. melanogaster*, and show that a cis-regulatory polymorphism associated with expression differences modulates lifespan variation in *Drosophila*. These results show that our experimental design allows for the estimation of the phenotypic effects of a single eQTL. The first report of *midway*’s involvement in *Drosophila* lifespan was recently confirmed using loss-of-function mutants (Pallares, Lea, et al., 2023). The changes in gene expression levels caused by the cis-eQTL that we validated here, contrasts the drastic changes in genome organization caused by complete loss-of-function. While complete loss-of-function mutants for midway show a significant decrease in lifespan (T50 = 33 for midway loss-of-function mutant and T50 = 47 for control populations) (Pallares, Lea, et al., 2023), here, we find that the cis-eQTL genotype that reduces midway expression actually increases lifespan. The contrast between those results highlights the benefit of validating genetic effects in a relevant genetic context: the biological insight obtained from lab-based experiments will be a more accurate representation of the effect of genetic variation in natural populations.

The broad applicability, simplicity, and low cost of the experimental approach we advance in this study offers a tractable system for validating allelic effects in diverse backgrounds and provides significant gain in statistical power to detect genotype-phenotype associations. While this study focuses on *D. melanogaster*, the general experimental paradigm we outline can be applied broadly to any model (or non-model) systems where focal individuals can be kept alive while being genotyped, and where controlled breeding is an option.

## Code and data availability

All code and data for reproducing the analysis presented here can be found on GitHub: github.com/ayroles-lab/SNPvalidation.

## Authors’ contributions

V.A., J.F.A., and L.F.P. designed the study. V.A. and L.F.P. performed the experiments. S.W., V.A., D.M., and L.F.P., performed the analysis. S.W., V.A., and D.M. wrote the original draft. S.W., V.A., D.M., J.F.A., and L.F.P. reviewed and edited the manuscript. All authors approved the manuscript.

## Acknowledgements

We thank all members of the Ayroles lab for their support.

